# Demonstration of RNA interference by bacterial feeding in *Stentor polymorphus*

**DOI:** 10.1101/843128

**Authors:** Mark M. Slabodnick

## Abstract

*Stentor* is a genus of large trumpet-shaped unicellular organisms in the ciliate phylum. Classically they have been used as models of cellular morphogenesis due to their large size and ability to regenerate, but some *Stentor* species have features that make them useful models for other types of studies as well. *Stentor polymorphus* is a widely distributed species that harbors green algal endosymbionts from the *Chlorella* genus. While interesting phenomenology in this species has been described, molecular tools have never been developed in this system. As technology has advanced, the use of emerging models like *S. polymorphus* has become more prevalent, and recently a set of transcriptomes for *S. polymorphus* was published. However, there are still technical hurdles to using *S. polymorphus* as an effective experimental system in the lab. Here I describe the identification and culture of a *S. polymorphus* population from North Carolina as well as the identification and cloning of homologs of α-tubulin and the morphogenesis gene *mob-1*. Additionally, I demonstrate that RNA interference (RNAi) by feeding is effective against both of these homologs in *S. polymorphus*. The phenotypes observed in *S. polymorphus* were similar to phenotypes previously validated in *S. coeruleus*, a related *Stentor* species. A direct comparison of feeding RNAi between the two species revealed that RNAi appeared to be less effective in *S. polymorphus*. The ability to perform RNAi in *S. polymorphus* strengthens its use as an emerging model for exploring mechanisms of unicellular morphogenesis and regeneration or host-symbiont interactions and suggests that RNAi by bacterial feeding might be more broadly effective across the *Stentor* genus.

## Introduction

The use of “non-model model organisms” or “emerging models” is becoming increasingly common in diverse fields of biological research facilitated by the rapid improvements in sequencing technologies and genetic techniques such as CRISPR and RNA interference (RNAi) which can be used to interrogate gene function [1, 2]. Some of these “new” systems are a return to organisms that were studied over a hundred years ago, but were never developed as experimental model systems. Many of these organisms exhibit fascinating biological phenomena and processes not apparent in the canonical set of model organisms, and thus can provide unique insights into biological mechanisms [1, 2]. For example, *Naegleria gruberii* is an ameboflagellate that moves by both amoeboid crawling and flagella-based swimming [3], and has led to insights about the evolution of cell motility mechanisms used across the tree of life [4, 5]. Because of their positions on the tree of life, volvox and choanoflagellates are two examples of models for the evolution of multicellularity, and both have yielded insights into how cells differentiate, stick together after division, and form organized multicellular structures [6-9]. This is by no means an exhaustive list, but represents some of the exciting work that is made possible by exploring these unique systems.

*Stentor polymorphus* is a large unicellular ciliate that is suited for development as an emerging model for the study of host-symbiont interactions as well as morphogenesis and regeneration. *Stentor* is a genus in the ciliate phylum that is renowned for its morphology and ability to regenerate. The most well-studied species in this genus, *Stentor coeruleus*, has recently been established as a model with tools such as a sequenced genome and ability to perform RNA interference (RNAi) by feeding [10, 11], but other *Stentor* species lack similar tools. *S. polymorphus* was first reported as a distinct species in 1773, although at the time it was called *Vorticella polymorpha* [12]. *S. polymorphus* cells are ∼250-500 um long and have a distinct green color [13] (Fig 1A) due to the presence of algal endosymbionts from the *Chlorella* genus (Fig 1B) [14]. Unlike many other *Stentor* species that possess colorful pigment granules, *S. polymorphus* lacks pigment except for the green color from the algal symbionts [13]. Similar to other *Stentor, S. polymorphus* are covered in cilia which are used for swimming and exhibit positive phototaxis [15]. They also have specialized cilia around their oral apparatus that are used for creating flow fields in the surrounding environment for capturing prey [16]. *S. polymorphus* forms clusters, containing dozens of individuals, that are anchored together onto plants, rocks, or pond debris (Fig 1C). Finally, *S. polymorphus* also possess the ability to rapidly and accurately regenerate after injury, which is the feature that made *Stentor* a focus of classical studies.

**Figure 1:**
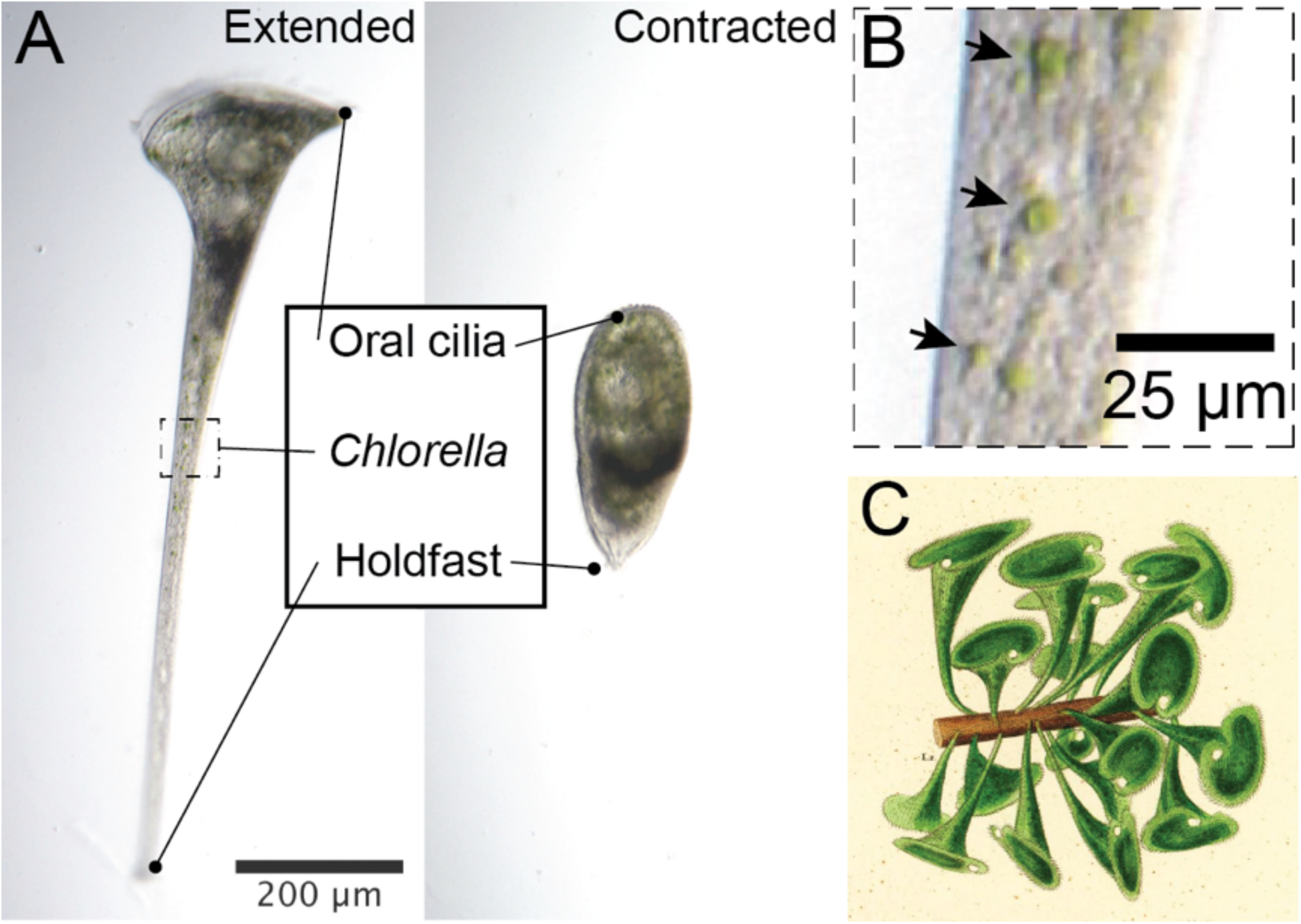
Stentor polymorphus. **(A)** An individual *S. polymorphus* cell imaged by brightfield microscopy. Cells are ∼250-500 μm in length with a trumpet shape and green *Chlorella* symbionts visible within the cell. **(B)** Inset from (A) showing *Chlorella* symbionts (arrows). **(C)** Colorized illustration from C. G. Ehrenberg [36] that depicts clustering behavior of *S. polymorphus* in natural samples.

*S. polymorphus* is an attractive candidate addition to existing host-symbiont interaction models like *Paramecium bursaria* and *Hydra viridissima*, as each of these organisms harbors related *Chlorella* symbionts [14, 17, 18]. Interestingly, an isolate of *S. polymorphus* has been identified which harbors a different, non-*Chlorella*, algal symbiont lacking a pyrenoid, an organelle used in carbon fixation [19]. How the reduced photosynthetic output of a symbiont effects the relationship to the host organism is an open question. Furthermore, a different isolate was recently identified that completely lacks algal endosymbionts [20]. How the reduced photosynthetic output of a symbiont effects the relationship with the host, and how a single species of *Stentor* can exhibit these different symbiont requirements is an open question. In addition to its potential as a model for host-symbiont interaction, like other *Stentor, S. polymorphus* would also be a useful model for morphogenesis and regeneration, and a recent study has published a set of transcriptomes of the regeneration process in *S. polymorphus* [21]. Not only did this study produce a set of *de novo* transcriptomes, but it also generated a large list of genes that are potentially involved with regeneration in *Stentor*. However, currently there are no published methods to interrogate gene function in *S. polymorphus*. Studying morphogenesis and regeneration in multiple unicellular systems could yield insights in conserved or diverse mechanisms that have developed throughout evolution.

Here I demonstrate that a method for RNAi by feeding is effective in an isolate of *S. polymorphus*. Targeting an α-tubulin homolog resulted in small rounded cells with diminished tails. Whereas targeting a *mob-1* homolog resulted in cells that lacked normal proportions and polarity. Additionally, the effectiveness of RNAi between *S. polymorphus* and *S. coeruleus* was directly compared. This study expands our toolkit of tractable methods to study gene function in an organism with the potential to reveal key insights into the biology of cell regeneration, symbioses, and other unique biology.

## Materials and Methods

### Cell isolation and culture

*Stentor polymorphus* cells were initially isolated from a pond at UNC-CH (35°54’25.4”N, 79°02’09.4”W) and subsequently cultured in the lab. *Stentor coeruleus* was obtained commercially (Carolina Biological Supply, Item # 131598) and subsequently cultured in the lab. *S. polymorphus* cells were obtained on a sunny afternoon by collecting pond water along with duckweed or other plant matter floating near the surface. These samples were agitated by vigorous shaking or stirring to dislodge anchored *Stentor* and water was then poured into petri dishes for visual observation. Even after vigorous agitation, cells were still often found in clusters anchored to duckweed or other debris. Cultures were maintained in dishes containing 50mL of sterile filtered water from the pond where the cells were isolated. Cells were fed twice a week with *Chlamydomonas reinhardtii* (strain cc125, kind gift from the Marshall lab) as has been done previously for *Stentor coeruleus* [10, 22] and kept at 22°C in ambient room light with indirect sunlight from windows in lab. In this way the cells were under loosely controlled light/dark cycles, as it was previously reported that cells divide at night [13]. Additionally, cultures were supplied with three, one centimeter long, coconut fibers as a fibrous substrate for the cells to anchor (SunGrow Coconut Fiber, Amazon.com, ASIN:B079KDS24T).

### Gene cloning and plasmid construction

Genes from *S. coeruleus* were used as a reference to identify homologs in the *S. polymorphus* transcriptome. Primers were designed to amplify full length gene sequences from *S. polymorphus* cDNA (S1 Table). Genes were cloned into a modified pPR-T4P vector (S1 Fig) [23], containing a ccdB site flanked by a multi-cloning site, SapI sites, and two opposing T7 sites for dsRNA production. This vector, pSRV, is designed for easy cloning of a gene of interest that has been amplified with SapI sites and the corresponding 3 nucleotide overhangs as part of the primer sequence, based on the SapTrap method with ccdB negative selection [24]. The sequence for the gene of interest then replaces the ccdB cassette, which serves as a negative selection against uncut or resealed vector. The pSRV plasmid will be made available on Addgene.

### Alignment and Phylogenetic Analysis

28S rDNA sequences from other species were obtained from NCBI (S2 Table) and aligned using CLC Genomics Workbench (Qiagen) with the following parameters: Gap open cost = 10.0, Gap extension cost = 1.0, End gap cost = As any other, Alignment mode = Very accurate (slow). Aligned sequences were then trimmed using Gblocks [25]. Neighbor-joining trees were constructed using CLC Genomics Workbench (Qiagen) using the Juke-Cantor distance measurement and 10,000 bootstrap replicates.

### RNAi method

*S. polymorphus* or *S. coeruleus* cells were washed and starved for ≥12 hours before the experiment began, with 25 cells per well containing 500 μL of filtered pond water. Experiments were successfully performed in both plastic 24-well flat bottom dishes (Olympus Plastics, Genesee Scientific, Cat. # 25-107) and 9-well spot plates (Corning Glass, VWR, Cat. # 89090-482). The HT115 strain of *E. coli* was used for the feeding vector [26], transformed with a plasmid containing the target gene flanked by two opposing T7 sites for double-stranded RNA production. HT115 cells were grown overnight in a 37°C shaking incubator under the selection of both Kanamycin and Tetracycline. Cultures were then diluted to OD600 = 0.1 and grown back to OD600 = 0.5 under the selection of only Kanamycin. Cultures were then induced to produce dsRNA by the addition of 1mM IPTG (Millipore Sigma, SKU. # I5502) and further incubated for 4 hours. Bacterial cells were then harvested by centrifugation and washed in filtered pond water. Bacterial cells were then aliquoted, centrifuged to remove media, and dry pellets were flash frozen in liquid nitrogen and stored at −80°C for up to one week. Aliquots contained 200 μL of the initial bacterial culture and were resuspended in filtered pond water daily and added to each well containing *Stentor* cells. To avoid buildup of detritus the *Stentor* were washed into a new well containing filtered pond water every other day. Cells were quantified, scored, and imaged during this wash step.

### Immunostaining

Cells were fixed and stained using previously published methods for *S. coeruleus* [10]. Briefly, cells were first washed with sterile pond water and then isolated in a minimal volume. Cells were then fixed with ice cold methanol and incubated for at least 20 minutes at −20°C, rehydrated using a 1:1 methanol:phosphate buffered saline (PBS) solution and incubated for 5 minutes at room temperature, then finally washed with 1x PBS for 10 minutes at room temperature. Cells were blocked in 1x PBS with 0.1% Triton-X-100 and 2% BSA (blocking buffer) and incubated for 1 hour at room temp. Cells were stained in suspension and allowed to settle to the bottom of the tube. Antibodies were diluted in blocking buffer, and mouse monoclonal anti-tubulin (clone DM1A, MilliPore Sigma, Cat. #T6199) was used at a 1:500 dilution for a primary antibody incubation of 1 hour at room temperature. After washing 3x in PBS, Alexa-488 goat-anti-mouse secondary antibody (ThermoFisher, #A-11017) was used at a 1:1000 dilution and incubated for 1 hour at room temperature. A final wash, 3x in PBS, was performed before mounting cells on a slide for imaging.

### Imaging

*Stentor* were imaged on a Zeiss Axiozoom V.16 equipped with PlanNeoFluar Z 1x/0.25 FWD 56mm and 2.3x/0.57 FWD 10.6mm lenses (Zeiss). Images were collected using a Canon EOS Rebel T6 camera (Canon) mounted to the Axiozoom using a T2-T2 1.6x SLR mount (Zeiss). Images were collected using the Canon EOS 3 Software (Canon). Images were adjusted and analyzed using ImageJ/Fiji [27]. Fluorescence images of immunostained *Stentor* were collected on a Nikon Eclipse Ti Spinning-disk confocal microscope (CSU-X1 spinning-disk head; Yokogawa, Tokyo Japan) using a Hamamatsu ImagEM X2 EM-CCD camera (C9100-13) and a 10x/0.30 numerical aperture (NA) Plan Fluor objective (Nikon, Tokyo, Japan).

### Data Analysis and Statistics

Data from RNAi experiments represents the mean values from three biological replicates. Phenotypes were qualitatively scored and data was recorded and analyzed using Microsoft Excel (Microsoft, Redmond, Washington), and error bars represent 95% confidence intervals.

## Results

### Isolation, Identification, and Culture of Stentor polymorphus

*Stentor polymorphus* was obtained from a pond on the campus of the University of North Carolina at Chapel Hill. Cells were first isolated by hand from pond water samples under a dissecting microscope, and putatively identified based on physical characteristics [13, 28]: 250-500 μm in length, trumpet shaped, presence of symbiotic green algae, and a moniliform shaped macronucleus (Fig 2A, 2B). The identity of *S. polymorphus* was next verified by 28S rDNA sequencing using universal primers to amplify gene sequences from isolated genomic DNA and compared against known sequences, as has been done previously to identify *Stentor* [20] (Fig 2C).

**Figure 2:**
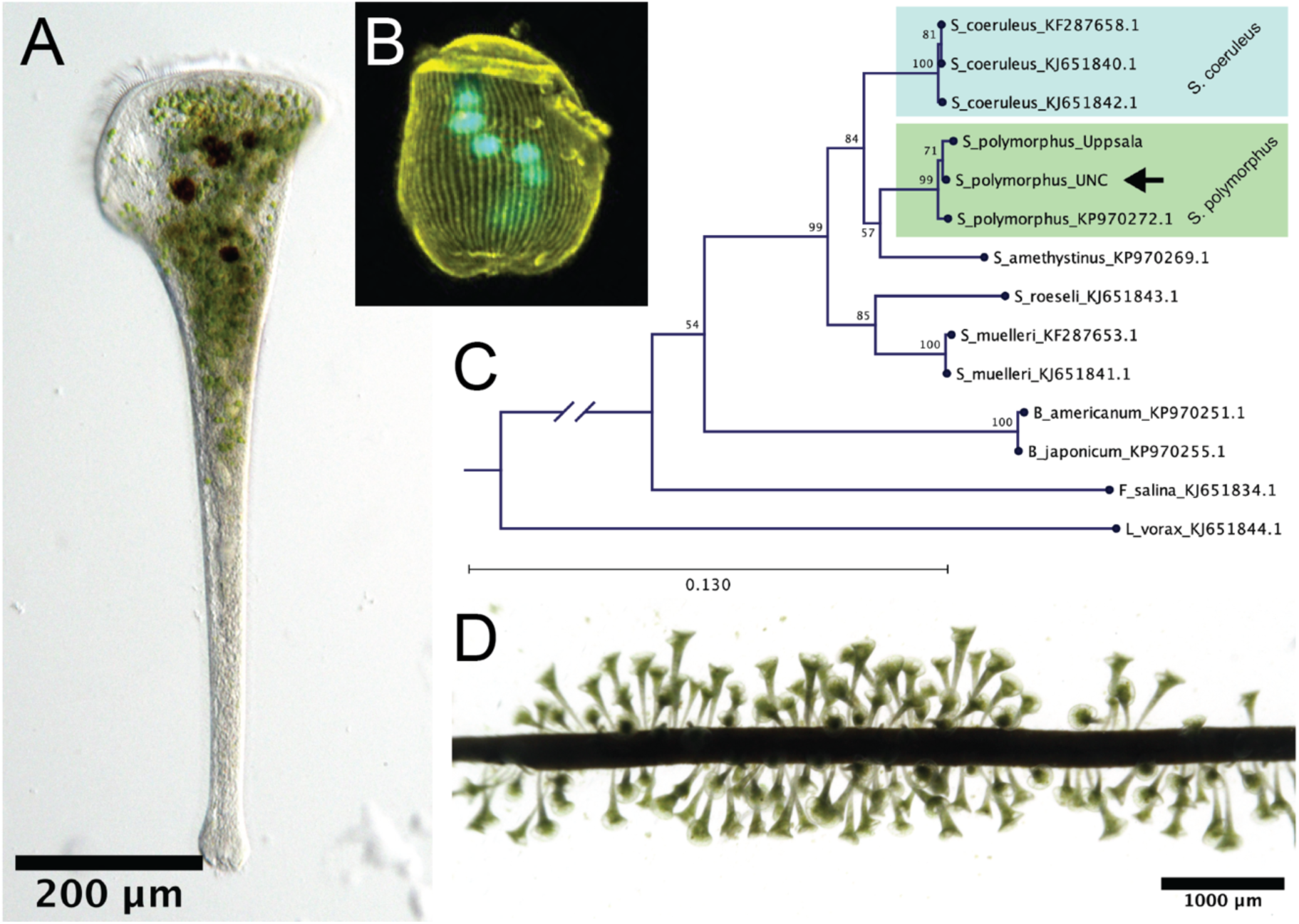
Isolation and culture of *S. polymorphus*. **(A)** Figure showing *S. polymorphus* isolate. **(B)** Immunostaining of *S. polymorphus* cell stained with anti-α-tubulin (yellow) and DAPI (cyan) showing the moniliform morphology of the nucleus. **(C)** Phylogenetic tree constructed using 28S rDNA sequences from selected ciliates to demonstrate the identity of the *S. polymorphus* isolate. *S. coeruleus* and *S. polymorphus* groups are indicated and *Loxodes vorax* was used as an outgroup. **(D)** *S. polymorphus* anchoring on coconut fibers in culture (compare to Fig 1C). When provided with coconut fibers, cells clustered on the fibers as shown and generally did not anchor elsewhere on the glass dish.

To maintain these organisms long-term in the lab, I established stable cultures of isolated *S. polymorphus* using the same laboratory culturing methods already used for *S. coeruleus* [22]. Briefly, *S. polymorphus* cells were washed in sterile filtered pond water (hereafter referred to as pond water) and ∼300 cells were placed in a small glass jar containing 50 mL of pond water. *Chlamydomonas reinhardtii* was cultured separately to be used as food, and added to the *Stentor* culture twice per week. Cultures were stored at room temperature on a lab bench and received natural light/dark cycles from a nearby window. Previous reports suggest that *S. polymorphus* divides at night [13], however dividing cells were observed during the day. One issue I found in culturing *S. polymorphus* was that cells would form large clusters in hard to reach areas of the culture dish, either in the bottom corners or at the top edge of the dish near the air-water interface. Having cells in these regions made it difficult to assess culture health and also led to accidental loss of cells at the air-water interface if the dish was disturbed. Therefore, I modified the previously published culture method and provided cells with a commercially available plant-based substrate, coconut fibers, on which to anchor. Cells readily anchored to these fibers and formed large clusters on small clippings that settled at the bottom of the dish (Fig 2D). This isolate has been kept in culture for over 12 months using the methods described.

### Gene Identification and RNA interference methods

Next, I sought to determine whether methods for RNA interference (RNAi) by bacterial feeding (referred to as RNAi feeding) were effective in *S. polymorphus*. I chose to test RNAi feeding in *S. polymorphus* using two genes, α-tubulin (*tba-1*) and *mob-1*, that had been previously shown to result in characteristic morphological phenotypes in *S. coeruleus* [10], I decided to try these same methods in *S. polymorphus*. In *Stentor*, tubulin is a critical component of the cytoskeleton that gives cells their shape, and *mob-1* is a component of a signaling pathway that is required for proper growth and patterning in ciliates [10, 29-31]. Therefore, I had the reasonable expectation that knockdown of these genes in *S. polymorphus* would result in similar quantifiable phenotypes, and I pursued both of these genes as standards for determining the effectiveness of RNAi in *S. polymorphus*.

The *S. polymorphus* homologs for the *tba-1* and *mob-1* genes were identified by reciprocal best hit BLAST from the published *S. polymorphus* transcriptome using the *S. coeruleus* protein sequences as a reference and cloned into an RNAi feeding vector (S1 Fig, S1 Table). The *tba-1* and *mob-1* transcripts chosen were 87.95/100 and 81.75/99 percent identical to their corresponding *S. coeruleus* nucleotide/protein sequences, respectively (S2A Fig). After cloning and sequencing the genes from the UNC isolate of *S. polymorphus*, I found that nucleotide sequences of *tba-1* and *mob-1* from *S. polymorphus* isolated at UNC were 94.46 and 97.84 percent identical, respectively, to the nucleotide sequences reported in the transcriptome generated from *S. polymorphus* cells isolated in Sweden (S2A Fig). All of the differences between each pair of sequences resulted in synonymous substitutions, and each pair of predicted protein sequences was 100% identical (S2B Fig). Therefore, despite the large geographical distance between these isolates, the differences between these two gene sequences were quite small.

Vectors for expressing dsRNA products were previously used for bacterial feeding in *S. coeruleus* [10]. I made modifications to the previously used vector that make cloning easier. These modifications were inspired by Golden Gate Assembly and SapTrap cloning methods [24, 32], and takes advantage of the Type IIS restriction endonuclease SapI (S1A Fig). As a Type IIS restriction enzyme, SapI cleaves outside of its recognition site allowing for the production of different *5’* overhangs with a single enzyme. Additionally, I inserted a ccdB cassette between the T7 promoters for negative selection against uncut and resealed vector backbone. With this design, SapI sites are added via PCR during amplification of the target gene, and SapI sites on either side of the ccdB cassette are used to produce corresponding *5’* overhangs in both the vector backbone and the gene insert (S1B Fig). Because the insertion of the target gene sequence results in a product lacking SapI recognition sites, digestion and ligation can be performed in the same tube at the same time.

Feeding of both *tba-1* and *mob-1* dsRNA-producing bacterial vectors resulted in specific morphological phenotypes similar to those previously observed in *S. coeruleus* [10]. Compared to control, cells fed bacteria expressing dsRNA targeting *tba-1* became small and round with shorter and thinner tails (Fig 3A, 3B). Cells fed bacteria expressing dsRNA targeting *mob-1* first became disproportioned (lost their stereotypical “trumpet” shape and became wider along their length, Disproportioned) (Fig 3C, left). Later, mob-1 knockdown cells began to develop multiple points that resembled posterior protrusions (Medusoid) (Fig 3C, right). While disproportioned *S. polymorphus* cells looked nearly identical to phenotypes previously observed in *S. coeruleus* [10], I observed that the posterior-like protrusions on medusoid *S. polymorphus* cells were not as extended, and the oral apparatus had a normal appearance. This might represent a difference in *mob-1* function between the two species, but might also represent a difference in RNAi efficiency.

**Figure 3:**
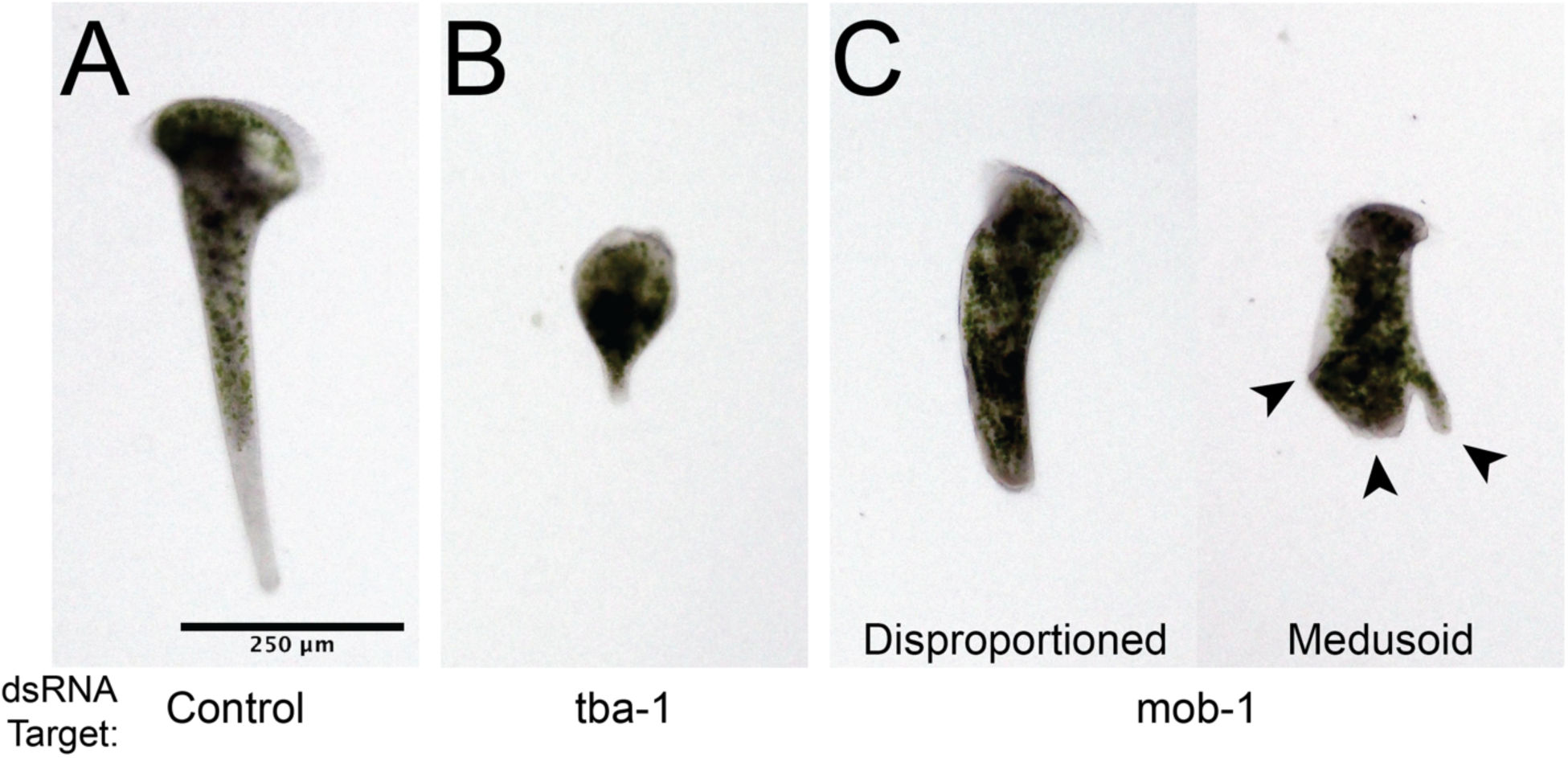
Targeting genes by RNA interference in *S. polymorphus* results in specific defects. **(A)** Targeting a control gene not present in *S. polymorphus* results in cells with a normal trumpet shape. **(B)** Targeting tba-1 in *S. polymorphus* results in small cells. **(C)** Targeting mob-1 in *S. polymorphus* results in two different phenotypes. Disproportioned cells (left) have an elongated central region. Medusoid cells (right) possess multiple posterior-like protrusions (arrowheads). All of the observed phenotypes are consistent with the results of gene knockdown in *S. coeruleus* where homologs were targeted.

In contrast to α-/β-tubulin knockdown phenotypes in *S. coeruleus*, mob-1 knockdown is generally 100% penetrant, and phenotypes manifest relatively rapidly, reproducibly, and in a specific temporal pattern with the disproportioned cells appearing first in the population before transitioning into medusoid cells over the course of 4-6 days [10]. Although experiments performed in *S. polymorphus* resulted in phenotypes that were similar to those previously observed in *S. coeruleus*, I noticed that the appearance of these phenotypes took several days to manifest, longer than what was previously reported for *mob-1* knockdown phenotypes in *S. coeruleus*. In order to characterize the differences between RNAi in these two *Stentor* species, I directly compared the timing of the appearance of phenotypes between *S. polymorphus* and *S. coeruleus* treated in parallel (Fig 4A, 4B), which confirmed that the phenotypes took longer to manifest in *S. polymorphus* as compared to *S. coeruleus*. Also, as compared to *S. coeruleus, S. polymorphus* medusoid cells had a relatively normal looking anterior, and the posterior protrusions were less pronounced. Despite this difference between the two species, these data show that methods for RNAi by feeding are effective in *S. polymorphus*.

**Figure 4:**
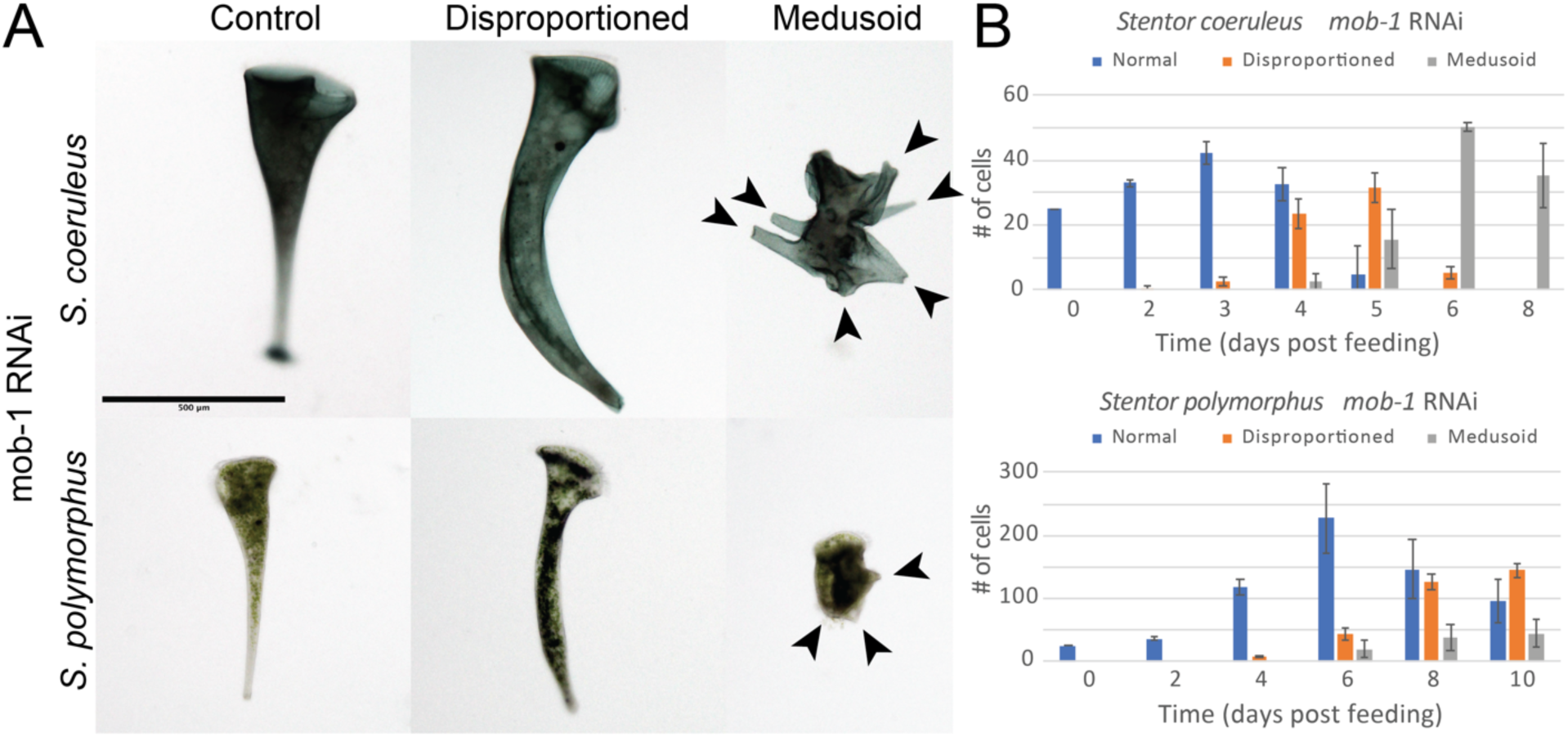
Comparison of mob-1 phenotype timing between *S. coeruleus* and *S. polymorphus*. **(A)** Images of Control (left) and mob-1 RNAi (right) cells comparing *S. coeruleus* (top) with *S. polymorphus* (bottom). Observed phenotypes are extremely similar between the two species, although medusoid cells retain a normal looking oral apparatus and the posterior protrusions (arrowheads) are not as elongated. Scale bar = 500 μm. **(B)** Data showing the appearance of phenotypes after feeding bacterial vectors targeting mob-1 in populations of either *S. coeruleus* (top) or *S. polymorphus* (bottom) cells over the course of the experiment. Error bars represent 95% confidence intervals.

## Discussion

Here I report that both culture and RNAi methods that are used for *Stentor coeruleus* are applicable to the closely related *Stentor polymorphus*. These results are of value because both of these organisms have been used previously for the study of single-cell regeneration, and both organisms have published transcriptomes of gene expression during the regeneration process. However, before this report there were only published methods for studying gene function in *S. coeruleus*. Here I have shown that RNAi by feeding in *S. polymorphus* is effective at producing nearly identical phenotypes to those observed in *S. coeruleus*, though with different timing. With the ability to culture *S. polymorphus*, identify and clone genes, and test gene function in this system, *S. polymorphus* can be used as a model for evolutionary comparisons of cell regeneration, host-symbiont interaction, or as a model for other fascinating biology.

Why RNAi was less effective in *S. polymorphus* is unknown and was not specifically probed in this study. Cells appear to consume similar amounts of food relative to their size, although it is possible that *S. coeruleus* is able to gather and process food faster than *S. polymorphus*. The reasons for this discrepancy in RNAi efficiency could be due to various factors. For example, *S. polymorphus* divides faster than *S. coeruleus* and so it could be due to difference in the cell cycles of these two species. Small RNAs are also known for playing critical roles in ciliate development and gene regulation [33-35], so these cell cycle differences might mean that the cell’s small RNA processing machinery is less available for RNAi. Or maybe the RNAi machinery in *S. coeruleus* is more highly expressed, more efficient, or otherwise more effective than in *S. polymorphus*. In any case, more genes will need to be tested to determine if this trend is true for more than just *mob-1*.

With the ability to perform RNAi now paired with the data from the published transcriptomic study of genes that are upregulated during regeneration in *S. polymorphus*, experiments can now be performed to test gene function and determine genes that are required for regeneration in both *S. coeruleus* and *S. polymorphus*. Additionally, the demonstration that RNAi is effective in *S. polymorphus* adds one more species that is amenable to study in the *Stentor* genus, and suggests that RNAi by feeding might be broadly effective across the *Stentor* genus and would be a valuable tool to study gene function in any sequenced species that can be cultured. Several other *Stentor* species can be easily identified and isolated from the wild, often from the same ponds, including: *S. igneus, S. roeselli, S. introversus, S. pyriformis, S. multiformis*, and *S. amethystinus*. Unfortunately, not all of these species have been able to be stably cultured in the lab and transcriptomic or genomic sequences are not yet available for any of them. However, with the decreased costs and increased accessibility of genome/transcriptome sequencing, it is exciting to consider the possibilities once these other *Stentor* species are pursued as emerging models.

## Acknowledgements

I thank members of the Goldstein lab, Tatyana Makushok, and Jennifer Heppert for helpful discussion and comments on the manuscript. This work was supported by the National Science Foundation Award IOS 1557432.

