## Supplementary material for "Demonstration of RNA interference by bacterial feeding in *Stentor polymorphus*": S1 Fig

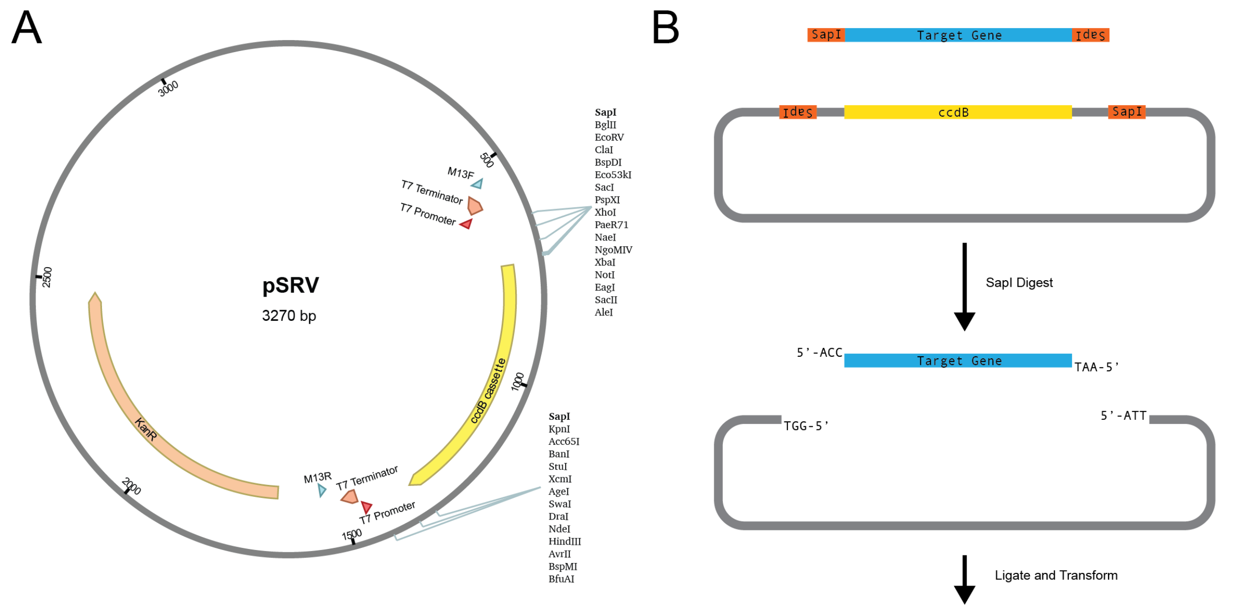


**<u>Supplemental Figure 1</u>: Map of pSRV RNAi feeding vector**. **(A)** Modified pPR-T4P vector compatible with SapI restriction cloning. A ccdB cassette is used for negative selection of uncut and resealed backbone. **(B)** The target gene of interest is PCR amplified with primer adaptors that add flanking SapI cut sites. SapI cut sites also flank the insertion site which produce compatible sticky ends after restriction digest as shown. Because only the desired product lacks SapI sites, digestion and ligation can be performed in the same tube for increased convenience and efficiency.
