## Supplementary material for "Demonstration of RNA interference by bacterial feeding in *Stentor polymorphus*": S2 Fig

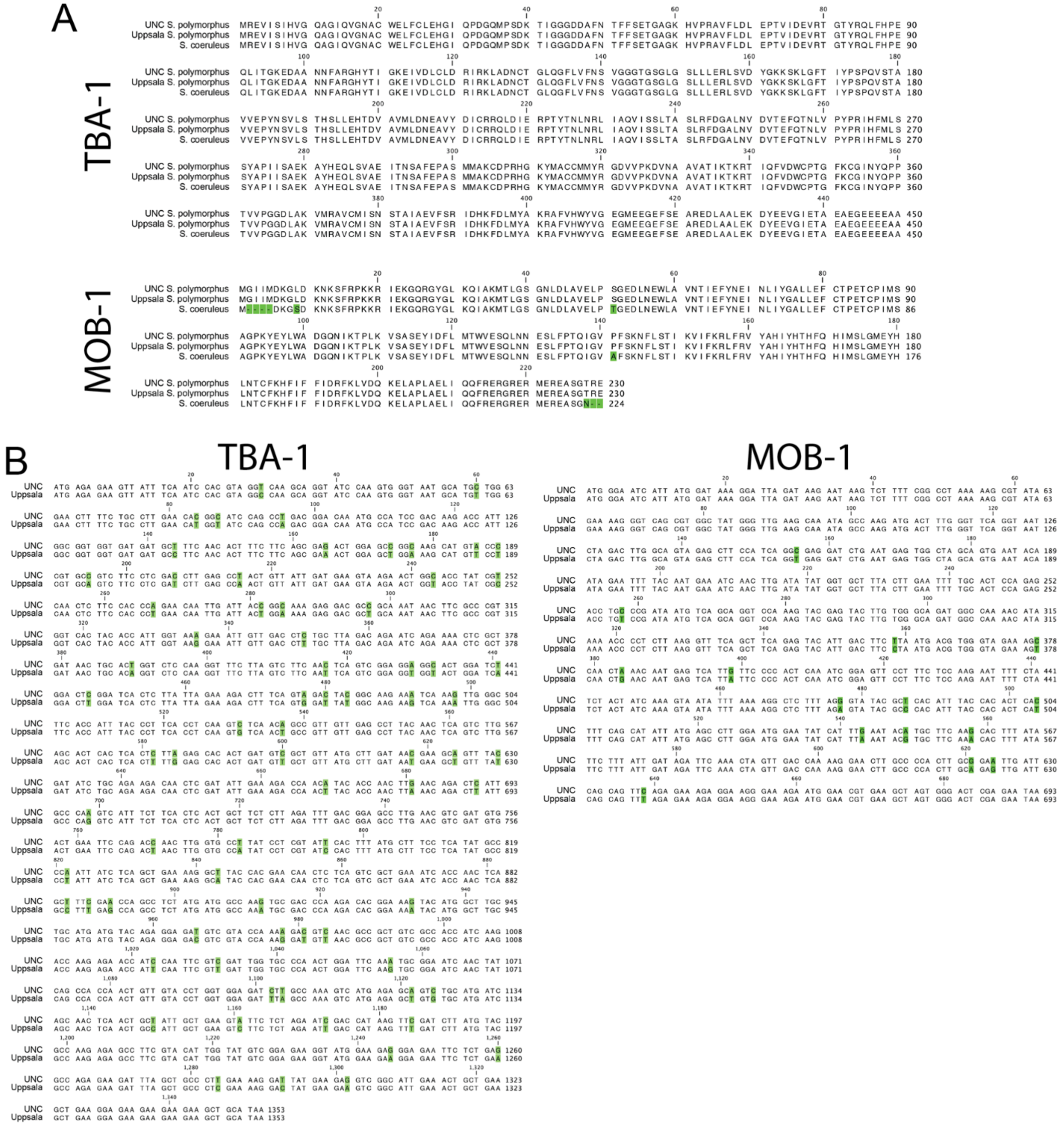

**<u>Supplemental Figure 2</u>: Identification and comparison of genes from *S. polymorphus* isolates (A)** Alignment of TBA-1 (top) and MOB-1 (bottom) protein sequences from the published *S. coeruleus* genome and *S. polymorphus* transcriptome [21] as compared to the UNC *S. polymorphus* isolate, differences are highlighted **(B)** Nucleotide alignments between published (Uppsala) and isolated (UNC) sequence shown in codon groupings, differences are highlighted. None of the SNPs that occur between the two isolates cause amino acid changes in the predicted protein sequence.
