## Supplementary material for "Demonstration of RNA interference by bacterial feeding in *Stentor polymorphus*": S1 Table

| Gene | *mob-1* | *tba-1* |
|---|---|---|
| *Query: S. coeruleus* gene ID | SteCoe_20658 | SteCoe_14841 |
| *RBBH: S. polymorphus* transcript ID | SP19394.1 | SP31362.593 |
| Forward Primer | atccatggGCTCTTCGACC**ATGGGAATCATTATGGATAAAGGATTAGATAAG** | atccatggGCTCTTCGACC**ATGAGAGAAGTTATTTCAATCCACGTAGG** |
| Reverse Primer | gcaggcctGCTCTTCGAAT**TTATTCTCGAGTCCCACTAGCTTCAC** | gcaggcctGCTCTTCGAAT**TTATGCAGCTTCTTCTTCTTCTCCTTC** |

**Supplemental Table 1: Identification and cloning of *mob-1* and *tba-1* homologs in S. polymorphus.** Gene IDs from the *Stentor coeruleus* genome browser (stentor.ciliate.org) used to identify reciprocal best BLAST hits (RBBH) from the *Stentor polymorphus* transcriptome [21]. Gene specific regions of the primers are bold, the rest of the primer sequence was used for cloning into pSRV.
