## Supplementary material for "Demonstration of RNA interference by bacterial feeding in *Stentor polymorphus*": S2 Table

| GenBank Accession Number | Genus | species |
|---|---|---|
| KJ651843.1 | *Stentor* | *roeseli* |
| KJ651841.1 | *Stentor* | *muelleri* |
| KF287653.1 | *Stentor* | *muelleri* |
| KJ651840.1 | *Stentor* | *coeruleus* |
| KJ651842.1 | *Stentor* | *coeruleus* |
| KF287658.1 | *Stentor* | *coeruleus* |
| KP970272.1 | *Stentor* | *polymorphus* |
| KP970269.1 | *Stentor* | *amethystinus* |
| KP970255.1 | *Blepharisma* | *japonicum* |
| KP970251.1 | *Blepharisma* | *americanum* |
| KJ651834.1 | *Fabrea* | *salina* |
| KJ651844.1 | *Loxodes* | *vorax* |

**Supplemental Table 2: 28S rDNA sequences.** List of GenBank accession numbers for reported 28S rDNA sequences used in the phylogenetic analysis along with the corresponding genus and species for each sequence. The 28S sequence from the previously reported *S. polymorphus* transcriptome that was used in this analysis was: SP31362.262
